# Imaging-induced stress causes divergent phase transitions of RNA binding proteins in the *C. elegans* germ line

**DOI:** 10.1101/2021.10.17.464158

**Authors:** Mohamed T. Elaswad, Chloe Munderloh, Brooklynne M. Watkins, Katherine G. Sharp, Elizabeth Breton, Jennifer A. Schisa

## Abstract

One emerging paradigm of cellular organization of RNA and RNA binding proteins is the formation of membraneless organelles (MLOs). Examples of MLOs include several types of ribonucleoprotein granules that form via phase separation. A variety of intracellular pH changes and post-translational modifications, as well as extracellular stresses can stimulate the condensation of proteins into granules. For example, the assembly of stress granules induced by oxidative stress, osmotic stress, and heat stress has been well-characterized in a variety of somatic cell types. In the germ line, similar stress-induced condensation of proteins occurs; however, less is known about the role of phase separation during gamete production. Researchers who study phase transitions often make use of fluorescent reporters to study the dynamics of RNA binding proteins during live-cell imaging. In this report, we demonstrate that certain conditions of live-imaging *C. elegans* can cause an inadvertent stress and trigger phase transitions of RNA binding proteins. We show that imaging stress stimulates decondensation of multiple germ granule proteins, and condensation of several P-body proteins. Proteins within larger RNP granules in meiotically-arrested oocytes do not appear to be as sensitive to imaging stress as proteins in diakinesis oocytes of young hermaphrodites, with the exception of the germ granule protein PGL-1. Our results have important methodological implications for all researchers using live-cell imaging techniques. The data also suggest that the RNA binding proteins within large RNP granules of arrested oocytes may have distinct phases which we characterize in our companion paper.

## INTRODUCTION

Phase separation is an emerging principle of cellular organization. Among the well-studied membraneless organelles that assemble via liquid-liquid phase separation are ribonucleoprotein (RNP) granules composed of mRNA and RNA binding proteins. Types of RNP granules include processing bodies (P-bodies) and stress granules in an array of cell types, and germ granules in the germ line (Corbet and Parker 2019). Many RNA binding proteins in RNP granules can undergo reversible phase transitions among diffuse, liquid, and gel-like phases. Transitions among these phases can function as an adaptation to environmental and intracellular changes, and regulate mRNA metabolism (Alberti and Carra 2018). In a variety of somatic cells, stress granules are induced by unfavorable environmental conditions such as oxidative stress, temperature changes, hypoxia, and osmotic stress to promote cellular homeostasis (Ivanov et al., 2019). Similarly, condensates can be induced by stress in the germ line; examples include stress granules in mouse spermatocytes, reticulated sponge bodies and large U-bodies in *Drosophila* egg chambers, and large, hybrid RNP granules in *C. elegans* oocytes (Schisa, 2019).

Proper regulation of phase transitions is critical, as dysregulation of RNP granules is associated with disease states such as cancer, cardiovascular disease, and neurodegenerative diseases (Wang et al., 2022). For example, in ALS mutations, the FUS protein phase separates into abnormal neuronal granules, and ectopic aggregates of FMRpolyG protein in ovarian stromal cells are associated with Fragile-X-associated primary ovarian insufficiency (Ganion 1991; Buijsen et al. 2016; Friedman-Gohas et al. 2020). While a growing number of *in vivo* studies started to build on *in vitro* observations that demonstrate a role for multivalent interactions in driving the condensation and decondensation of proteins, the precise regulation and function of phase transitions in oogenesis is not well understood.

The *Caenorhabditis* germ line provides an excellent model to study questions of RNA binding protein condensation (Schisa et al. 2001; Jud et al. 2008; Wood et al. 2016). In a young *C. elegans* hermaphrodite, oocytes undergo meiotic maturation every 23 minutes; however, within a few days of adulthood, the sperm become depleted, and meiosis arrests an extended time (McCarter et al. 1999a). In the arrested oocytes, several RNA binding proteins condense into large RNP granules that are up to 20 times larger than a typical cytoplasmic P granule in oocytes (Schisa et al. 2001). The condensation is reversible if sperm are resupplied via mating, or major sperm protein is injected (Schisa et al. 2001; Jud et al. 2008). Several mRNAs and diverse types of RNA binding proteins are detected in the large RNP granules including P-granule proteins, P-body proteins, stress granule proteins, and other RNA binding proteins associated with translational regulation such as the KH-domain protein MEX-3 and PUF domain protein PUF-5 (Schisa et al. 2001, Jud et al. 2008; Noble et al. 2008; Hubstenberger et al. 2013). Based on their composition, the large, hybrid RNP granules are hypothesized to regulate mRNA metabolism during extended delays in the fertilization of oocytes.

In this study, we uncover a live imaging-induced stress that causes divergent phase transitions of RNA binding proteins in oocytes. While PGL-1 and GLH-1 decondense in diakinesis oocytes of young hermaphrodites in response to imaging stress, MEX-3, CGH-1, and CAR-1 undergo condensation into granules. We show that these proteins are generally not as sensitive to imaging stress when they are already condensed into large RNP granules in arrested oocytes. The differential stress responses may suggest distinct phases of RNA binding proteins within large RNP granules. Finally, our results have important implications for all researchers using live cell-imaging methods, especially those studying phase transitions.

## METHODS

### Strains and maintenance

All worms were grown on nematode growth media using standard conditions at 20°C (Brenner 1974) unless specified. Strains used include: DG4269 mex-3(tn1753[gfp::3xflag::mex3]), OD61 ItIs41[pAA5; pie-1::GFP-TEV-Stag::CAR-1; unc-119(+)], JH3644 *fog-2(g71)* V; meg-3(ax4320)[meg-3::mCherry)]X, CL2070 dvIs70 [hsp-16.2::GFP + rol-6(su1006)], SJ4005 zcIs4 [hsp-4::GFP] V, LD1171 IdIs4 [gcs-1p::GFP + rol-6(su1006)], OH16024 daf-16(ot971[daf-16::GFP]), JH3269 pgl-1(ax3122[pgl-1::gfp]), DUP64 *glh-1(sam24[glh-1::GFP::3xFLAG])*, JH1985 *unc-119(ed3); axIs1436[pCG33* pie-1prom:LAP::CGH-1]. Some strains were crossed into CB4108 *fog-2(q71)* as noted. Worms were synchronized using the hypochlorite bleach method.

### Microscopy and Image analysis

Worms were picked onto slides made with 2% agarose pads and paralyzed using 6.25mM levamisole. To image diakinesis oocytes of young hermaphrodites, synchronized 1 day post-L4-stage worms were used. To image meiotically arrested oocytes in *fog-2* strains, L4 females were separated from males and imaged 1 or 2dpL4 as indicated in figure legends.

Images were collected using a Leica compound fluorescence microscope (low magnification images of whole worms in Fig. 4) or a Nikon A1R laser scanning confocal microscope (all other images). All images for a given strain were collected using identical levels and settings. The number of granules in oocytes and total integrated density (intensity of fluorescence in granules) was determined using ImageJ particle analysis. In Figure 4, the threshold to categorize dispersed PGL-1 was determined by the lowest fluorescence intensity in granules seen in the control 0-10 min. worms. The threshold for condensed CAR-1 was determined to be >3x higher intensity than in control 0-10 min. worms.

### Imaging using oxidative stress and ER stress reporters

The positive control for *gcs-1*::GFP was incubating young adults at 34°C for 3.5 hrs and recovering at 20°C for 1 hr before imaging (An and Blackwell 2003). The positive control for *hsp-16*.*2*::GFP was 34°C for 2 hrs and recovering at 20°C for 12 hr before imaging (Strayer et al. 2003). The positive control for *hsp*-4::GFP was 8 hours at 30°C before imaging (Bischof et al. 2008). Controls were compared to worms imaged within <25 min. or 60-90 min. after being prepared. GFP levels in images were qualitatively scored.

### Statistical analysis

Sample sizes were determined using G*Power 3.1 for power analyses; all experiments were blinded and done at least in triplicate. Data are presented as mean ± SEM unless otherwise indicated. Statistical analyses were performed on GraphPad Prism 9.1, and specific tests are noted in figure legends. P-values <0.05 were considered statistically significant.

### Data Availability

All strains are available at the *Caenorhabditis* Genetics Center (CGC) or on request from the Schisa lab or other *C. elegans* labs. The authors affirm that all data necessary to confirm the conclusions in this article are included in the article, figures, and tables.

## RESULTS AND DISCUSSION

### Imaging conditions can inadvertently induce stress and phase transitions

The regulation of RNA binding protein phase transitions in the germ line is not well understood. While investigating candidate regulators of the RGG domain-protein PGL-1 in the adult germ line, we noticed unanticipated dispersal of PGL-1 during imaging of control RNAi-treated worms. Early in the imaging session, PGL-1 was detected in punctate P granules throughout the adult germ line as expected (Strome and Wood 1982); however, over extended time (referred to as late), PGL-1 became dispersed in oocytes and the distal core, appearing at increased levels throughout the cytosol and decreased levels in punctate granules (Fig. 1A). To determine if our imaging conditions were inadvertently inducing a stress response, we used the DAF-16::GFP reporter strain (Aghayeva et al. 2020). The DAF-16 FOXO (forkhead box class O) transcription factor translocates from the cytoplasm to the nucleus in response to unfavorable conditions such as heat stress, oxidative stress, starvation, or when cellular repair is needed (Oh et al. 2005; Henderson et al. 2006). We categorized the subcellular localization of DAF-16 as cytoplasmic, nuclear, or intermediate (mix of cytoplasmic and nuclear) (Oh et al. 2005). We found that DAF-16 was cytoplasmic or intermediate in the large majority of worms imaged within 25 minutes of slides being prepared (worms picked into 6.25 mM levamisole on a 2% agarose pad and coverslip added), and DAF-16 was nuclear in less than 10% of worms (Fig. 1B,C). For the purpose of a contrasting experiment we chose a much later time interval, imaging 60-90 minutes after slides were prepared. In contrast to the early-imaged worms, 73% of worms that were scored late had nuclear localization of DAF-16, and none were cytoplasmic. Based on these results, we consider extended imaging conditions as imaging stress.

**Figure 1.**
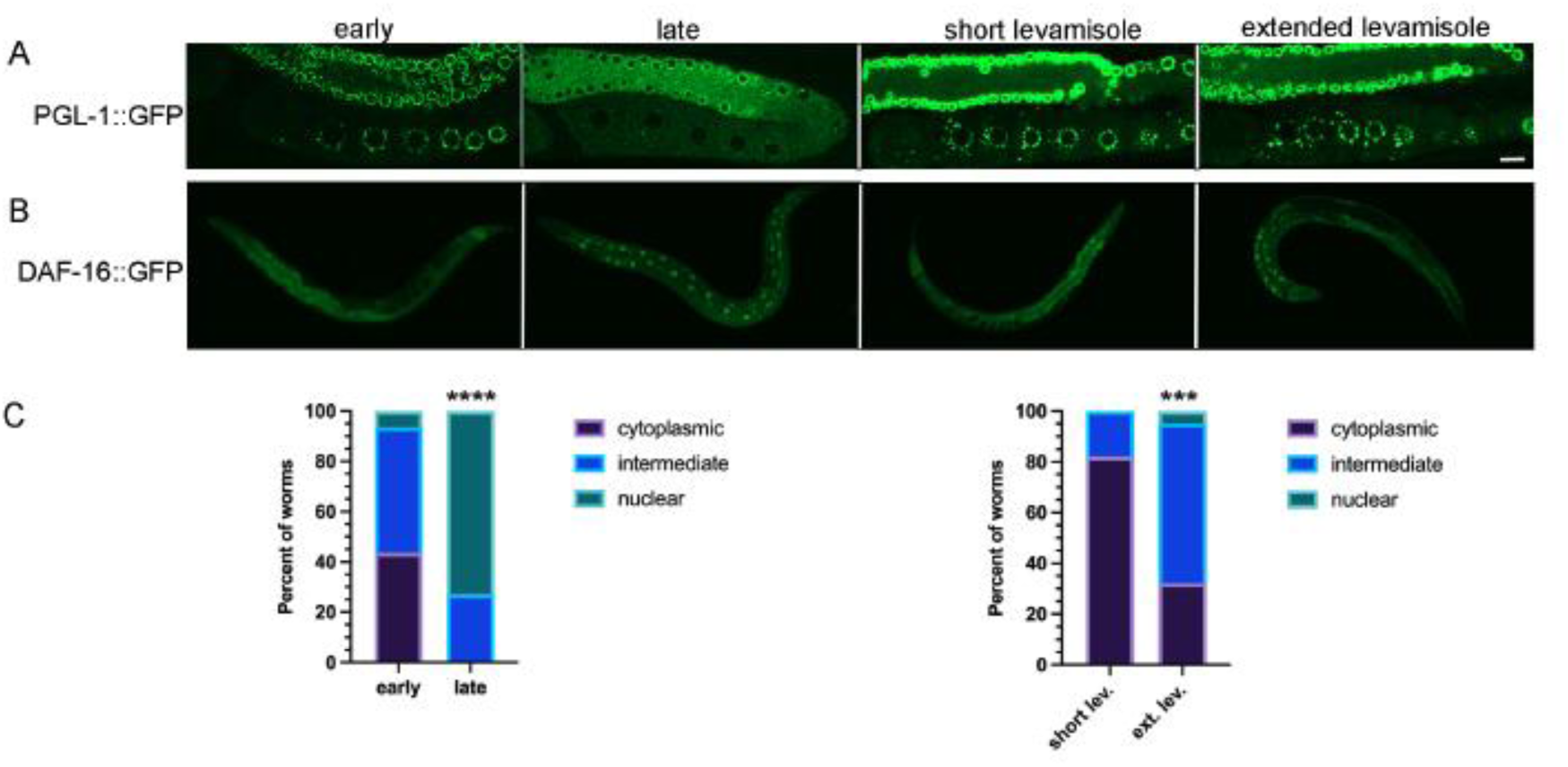
Imaging conditions can inadvertently induce stress. (A) Subcellular localization of PGL-1::GFP in the germ line: early during imaging process (within 25 min.) or after extended time imaging (60 min.); and after short (15 min.) or extended (60 min.) exposure to 6.25mM levamisole prior to mounting worms on a slide. Proximal oocytes are oriented on the bottom left in all images. Scale bar is 10 µm. (B) Subcellular localization of DAF-16::GFP in young hermaphrodites: imaging was done early (within 25 min.) or late (after 60-90 min.) of slides being prepared; and after short (15 min.) or extended (60 min.) exposure to 6.25mM levamisole. (C) Graphs show the percent of DAF-16::GFP worms with cytoplasmic distribution, intermediate distribution, or nuclear localization. Statistical significance was determined using the Fisher Exact Test. **** indicates p<0.0001; *** indicates p<0.001. n=15 (early), 16 (late), 28 (short), 19 (extended).

Levamisole is a paralytic agent frequently used as an anesthetic to mount worms for imaging. While one study showed that DAF-16 remains cytoplasmic after worms are exposed to 1mm levamisole and recovered for 30 minutes (Manjarrez and Mailler 2020), exposure to levamisole can affect autophagy and result in increased numbers of GFP::LGG-1 puncta in somatic cells (Zhang et al. 2015). We therefore asked if extended exposure to levamisole is sufficient to induce PGL-1 decondensation or DAF-16 translocation to nuclei. We incubated PGL-1::GFP worms in a drop of 50 µl of 6.25mM levamisole for 15 or 60 min in a humidity chamber, prior to placing them on an agarose pad and imaging within 15 minutes. We observed punctate PGL-1 granules in both samples, with no sign of the dispersed PGL-1 observed in late-imaged worms (Fig. 1A). In a large majority of DAF-16::GFP worms incubated a short time in levamisole we observed cytoplasmic DAF-16, with only 18% having an intermediate phenotype of partial nuclear translocation (Fig. 1B,C). In contrast, 63% of worms incubated an extended amount of time in levamisole had intermediate DAF-16 translocation, and in 5% of worms DAF-16 was nuclear. Because the ‘extended levamisole’ worms exhibited a weaker DAF-16 stress response than the late-imaged worms, we conclude levamisole contributes to stress during imaging, but is not the only source of stress. Our results suggest that while levamisole can contribute to a mild DAF-16 stress response, additional aspects of extended imaging underlie the decondensation of PGL-1 out of granules.

### Imaging stress causes divergent phase transitions of RNA binding proteins

To determine the extent to which imaging stress affects phase transitions of RNA binding proteins in diakinesis oocytes of young hermaphrodites, we first quantitated the effects on two P-granule proteins using the same conditions as for DAF-16. As expected for PGL-1, given our initial observations during RNAi experiments (Fig. 1), we observed significantly fewer granules of PGL-1::GFP in oocytes, and fewer granules in the distal nuclei, late during imaging (Fig. 2A,B). We also observed significant decondensation of GFP::GLH-1 in oocytes and the distal nuclei although the dispersal of protein out of granules was not as penetrant as with PGL-1 (Fig. 2A,B). We conclude imaging stress can induce the decondensation of multiple P-granule proteins.

**Figure 2.**
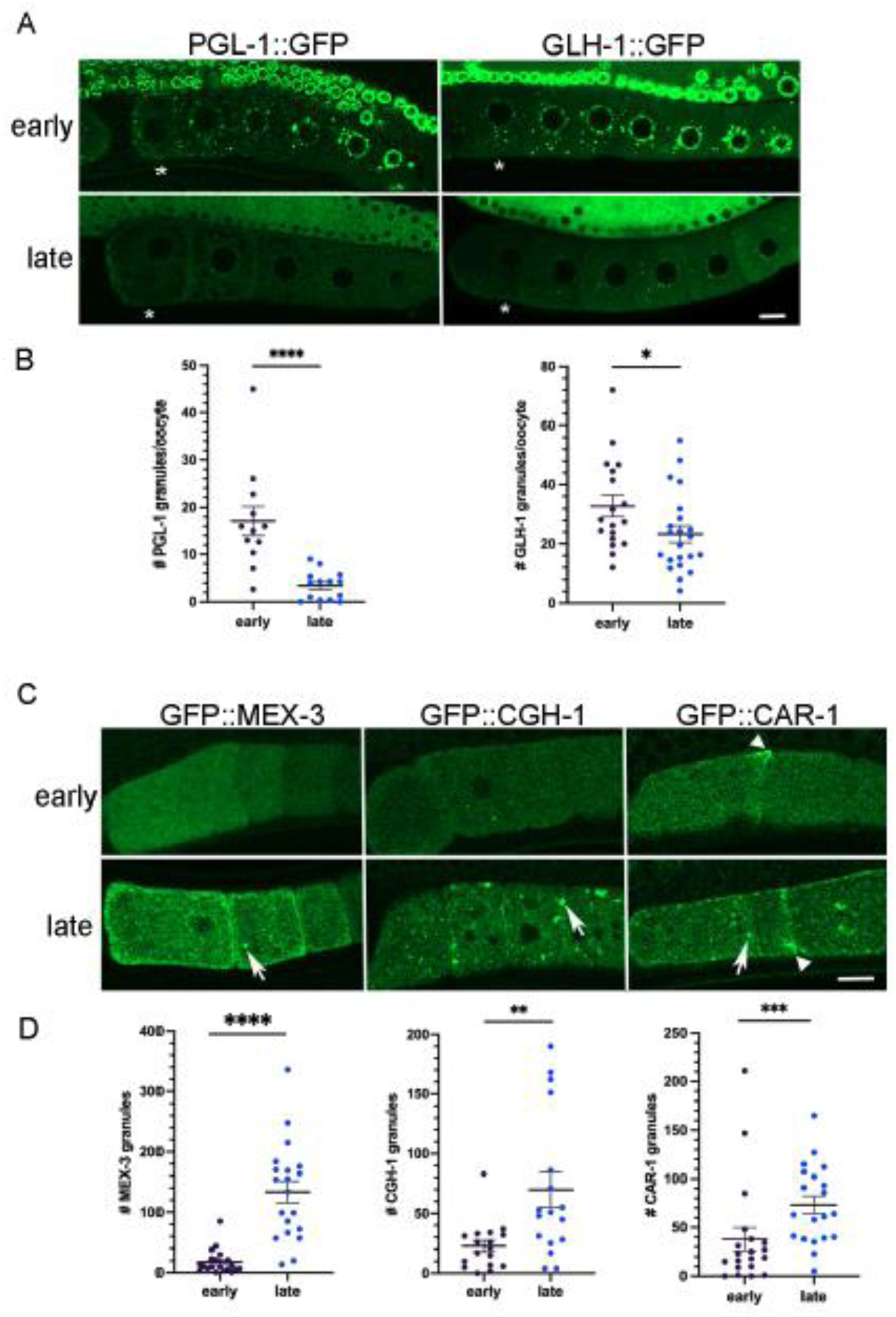
Imaging stress causes distinct phase transitions among RNA binding proteins. (A) Micrographs of GFP-tagged P granule protein reporter strains, PGL-1::GFP and GLH-1::GFP. Top row: Distribution of GFP in germ line early during imaging; Bottom row: Distribution of GFP in germ line after extended imaging (late) with strongest phenotypes shown. Asterisk marks the most proximal oocyte in each germ line. Smaller distal pachytene nuclei are visible at top of each image. Scale bar is 10 µm. (B) Graphs showing the number of GFP granules in a single Z-slice of proximal oocytes (see Methods). Statistical significance was determined using the Mann-Whitney test. **** indicates p<0.0001; * indicates p<0.05. n= 16-26. (C) Micrographs of GFP-tagged RNA binding protein reporter strains (MEX-3, CGH-1, and CAR-1) in most proximal oocytes. Top row: Distribution of GFP early during imaging; Bottom row: Distribution of GFP after extended imaging (late) with strongest phenotypes shown. Arrows indicate ectopic granules; arrowheads indicate cortical enrichment. Scale bar is 10 µm. (D) Graphs showing the number of GFP granules in a single Z-slice of proximal oocytes. Statistical significance was determined using the Mann-Whitney test. **** indicates p<0.0001, *** indicates p<0.001, ** indicates p<0.01. n=17-20. Error bars indicate mean ±SEM.

We next asked if imaging stress affects three RNA binding proteins that are normally dispersed throughout oocytes of young hermaphrodites. MEX-3 is a KH-domain protein that localizes to P granules in embryos, but is mostly diffuse throughout the cytosol of oocytes (Draper et al. 1996). In response to imaging stress the majority of GFP::MEX-3 protein appeared to remain decondensed throughout the cytoplasm; however, we detected a variable but significant increase in the number of small GFP::MEX-3 granules enriched cortically in oocytes (Fig. 2C,D). For phenotypes detected predominantly at the oocyte cortex, we chose to include cortical confocal slices that typically do not include the nuclei (Fig. 2C). CGH-1 and CAR-1 are orthologs of the P-body proteins Me31B/RCK and Trailerhitch/RAP55/Lsm14 (Schisa 2012). Both proteins are detected in P granules and throughout the cytosol of oocytes (Navarro et al. 2001; Boag et al. 2005). After imaging stress, the number of GFP::CGH-1 granules was variably increased; in a small number of worms, more than 7 times as many granules were detected compared to the control (Fig. 2C,D). Imaging stress also resulted in increased numbers of GFP::CAR-1 granules that were enriched at the nuclear envelope, as well as enriched along the cortical membrane (Fig. 2C,D). We also observed CAR-1 enrichment in non-spherical aggregates, distinct from the spherical granules, at the cortex in a subset of control and stressed worms (Fig. 1C arrowhead). Taken together, we conclude imaging stress can induce opposing RNA binding protein phase transitions in diakinesis oocytes of young hermaphrodites. MEX-3, CGH-1, and CAR-1 proteins undergo condensation into granules, opposite of the decondensation observed for PGL-1 and GLH-1.

The dynamics of proteins within granules can decrease as the size of RNP granules increases (Hubstenberger et al. 2013). Moreover, the properties and functions of RNA binding proteins can be disrupted upon phase transitions. For example, in *Drosophila* egg chambers in the absence of the E3 ubiquitin ligase HECW, the Me31B RNA binding protein transitions from small granules in a liquid phase to larger granules in a gel-like phase, and translational repression of target mRNAs is deregulated (Fajner et al. 2021). Since slower dynamics and increased stability can occur as RNP granules increase in size, we asked if imaging stress affects phase transitions of four RNA binding proteins that condense into large RNP granules in meiotically arrested oocytes of *fog-2* females. In early-imaged oocytes, we observed large cortical granules of GFP::MEX-3 and GFP::CGH-1 as expected (Schisa et al. 2001; Jud et al. 2008; Noble et al. 2008). After imaging stress, we did not detect any significant changes in the number of granules or the intensity of GFP fluorescence in granules, suggesting no major change in condensation (Fig. 3A,B). In contrast, PGL-1 granules were significantly reduced in number and intensity after imaging stress, compared to the ‘early’ control where a heterogeneous mix of small and very bright, large granules was detected as expected (Fig. 3A,B; (Jud et al. 2008; Noble et al. 2008)). Thus, we conclude imaging stress induces decondensation of PGL-1 out of large granules in arrested oocytes, similar to its effect on the small P granules in oocytes of young hermaphrodites. Lastly, we examined the effect of imaging stress on MEG-3. MEG-3 is an intrinsically disordered protein normally detected at low levels in oocytes and strongly localized to P granules in embryos (Wang et al. 2014). We found that during extended meiotic arrest in a *fog-2* background, MEG-3 underwent a dramatic phase transition and condensed into large granules in oocytes, as has been previously reported (Fig. 3A, Putnam et al. 2019). We detected no significant changes in MEG-3 granule number or intensity after extended imaging (Fig. 3A,B). Taken together, these data suggest that while MEX-3 and CGH-1 are sensitive to imaging stress in diakinesis oocytes and condense into granules, if the proteins are already condensed into large granules in arrested oocytes, they are more stable during imaging stress. Our data also suggest MEX-3, CGH-1, and MEG-3 phases of large RNP granules may be less liquid-like than PGL-1 which is sensitive to imaging stress in both diakinesis and arrested oocytes.

**Figure 3.**
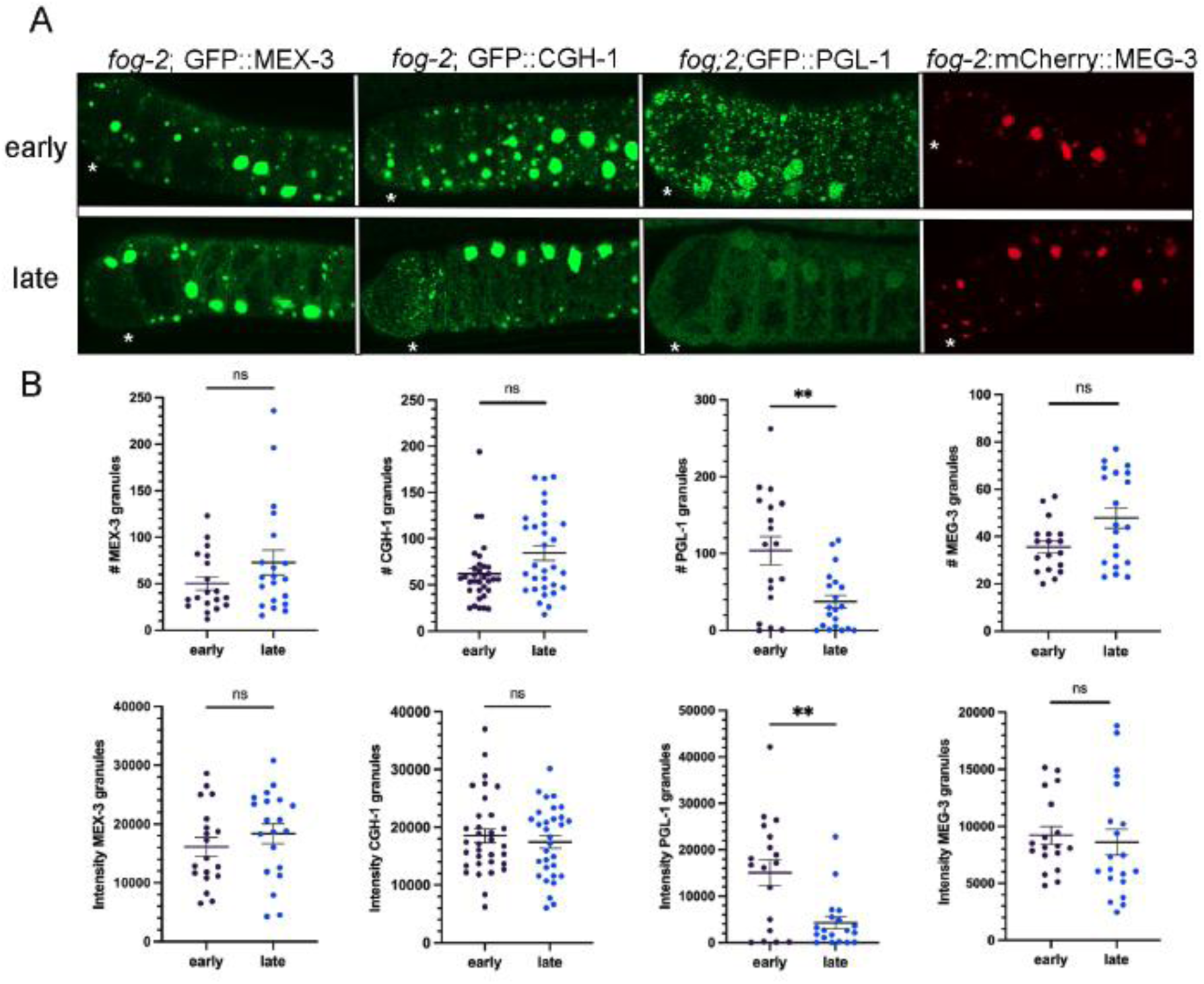
Imaging stress preferentially affects PGL-1 within large RNP granules of meiotically arrested oocytes. (A) Micrographs of GFP-tagged RNA binding proteins (MEX-3, CGH-1, PGL-1, MEG-3) in a *fog-2* background. Top row: Distribution of GFP in arrested oocytes early during imaging. Bottom row: Distribution of GFP in arrested oocytes after extended imaging (late). Asterisk marks the most proximal oocyte in each germ line. Scale bar is 10 µm. (B) Graphs showing either the number of GFP granules or the integrated density of GFP in granules in a single Z-slice of proximal oocytes (see Methods). Statistical significance was determined using the Mann-Whitney test. ** indicates p<0.01, ns indicates not significant. n=18-33. Error bars indicate mean ±SEM.

### Extended imaging does not appear to induce oxidative stress or ER stress

Once we established extended imaging can modulate phase transitions, we wanted to identify the type of stress induced by the imaging conditions. We used two reporter strains to ask if oxidative stress is induced: *gcs-1p*::gfp and *hsp-16*.*2*::gfp. GCS-1 (gamma-glutamyl-cysteine synthetase heavy gene) is a detoxification gene that is induced in the intestine in response to oxidative stress, including heat-induced increases in intracellular reactive oxygen species (ROS) (An and Blackwell 2003; Wang et al. 2010). HSP-16.2 is a small heat shock protein that is also induced by oxidative stress (Link et al. 1999). In the positive control, heat-stressed *gcs-1p*::gfp worms, we observed moderate or high GFP expression in 100% of the worms (Fig. 4A). In contrast, in oocytes assayed early during imaging without heat stress, moderate or high levels of GFP were detected in only 4% of worms, and the GFP expression did not significantly increase after imaging stress (Fig. 4B). In these experiments, levels of expression were assessed qualitatively using reference images. In the positive control, heat-stressed *hsp-16*.*2*::gfp worms, high levels of GFP expression were seen in 100% of worms. In contrast, low levels of expression were observed in worms imaged either early or after imaging stress (Fig. 4A,B). We also asked if ER stress is induced during imaging by using the *hsp-4*::gfp reporter. HSP-4 is the immunoglobulin heavy chain-binding protein (BiP) homolog, and a reporter for the unfolded protein response (UPR) that prepares cells for expansion of the ER (Shen et al. 2001; Kapulkin et al. 2005; Walter and Ron 2011). In the positive control, after 8 hours of 30°C heat stress, we observed moderate or high GFP levels in 96% of worms as expected (Bischof et al. 2008). In contrast, 100% of worms imaged early had low levels of GFP, and 98% of worms maintained low GFP levels after imaging stress (Fig. 4B). Overall, these results suggest the oxidative, ER stress, and heat stress pathways are not stimulated at high levels by imaging stress, and a different type of stress is induced by extended imaging. Prior studies report that a variety of stresses including osmotic stress, starvation, and anoxia can trigger the condensation of RNA binding proteins in the germ line (Jud et al. 2008; Huelgas-Morales et al. 2016; Davis et al. 2017). We speculate that a hypoxic stress may occur during extended imaging due to prolonged coverslip exposure; however, we were not able to find a suitable reporter to test this hypothesis in adults.

**Figure 4.**
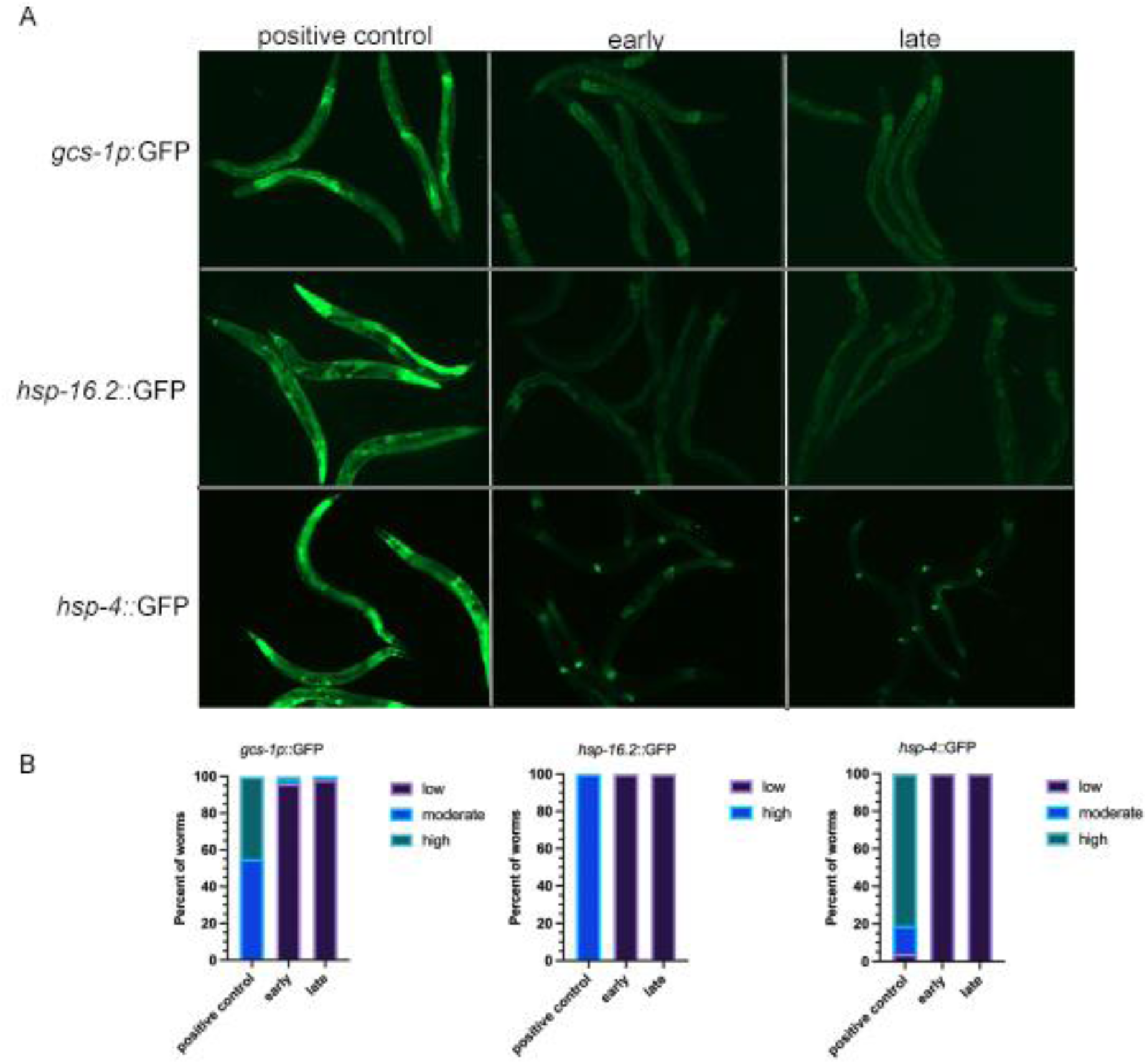
Extended imaging does not appear to induce oxidative or ER stress. (A) Micrographs of *gcs-1*::GFP, *hsp-16*.*2*::GFP, and *hsp-4*::GFP young adult worms either early during imaging or after imaging stress. The positive control for each was heat stress (see methods). (B) Graphs show the percent of worms with low, moderate, or high levels of expression of GFP. No significant differences were detected between early and late-imaged worms in any of the reporter strains. n=27-54.

### Imaging conditions can induce a rapid stress response

Having established that imaging stress can differentially trigger condensation or decondensation of RNA binding proteins, we asked how quickly the stress occurs while imaging. We first examined DAF-16::GFP worms that were prepared prior to imaging for a maximum of 10, 20, or 30 minutes. In the majority of 0-10-minute worms, DAF-16 was cytoplasmic, and none had strong nuclear localization (Fig. 5A). In contrast after 11-20 minutes, DAF-16 was nuclear in 69% of worms and intermediate in the remaining 31% of worms (Fig. 5A). In the 21-30 minute worms, DAF-16 was nuclear in >80% of worms. These time course data indicate that some level of stress occurs within 11-20 minutes of worms being prepared for imaging, and is relevant for all researchers using live-imaging methods to consider.

**Figure 5.**
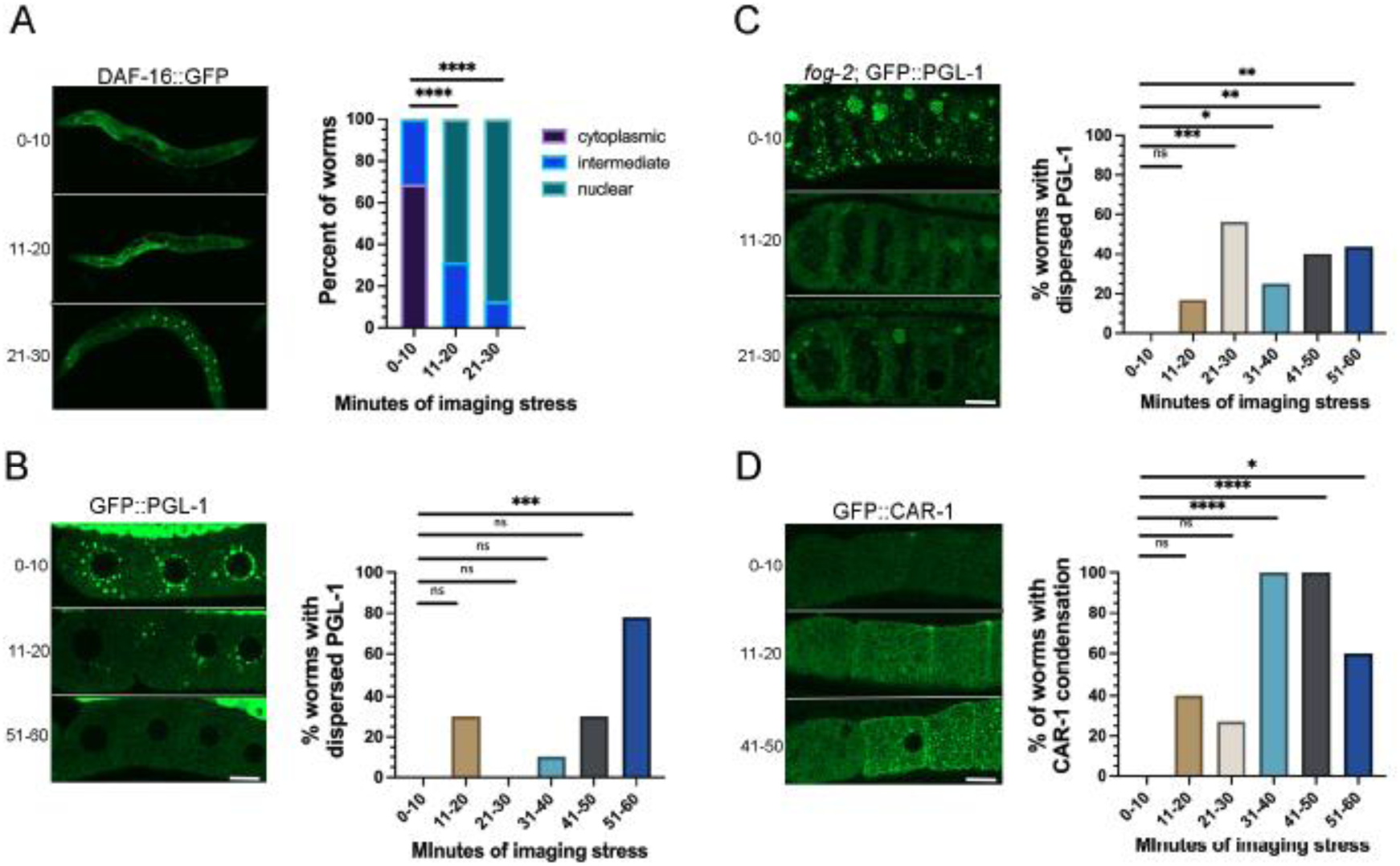
Imaging conditions can quickly trigger stress and modulate phase transitions. (A-D) Time course of the effects of imaging stress on DAF-16 nuclear translocation, and condensation of PGL-1 and CAR-1. For each reporter, micrographs of GFP-tagged strains are shown on the left with proximal oocytes oriented to the left, and graphs show the percent of worms with nuclear translocation (DAF-16) or phase transitions (PGL-1, CAR-1). Diakinesis oocytes in young hermaphrodites are shown in panels A, B, D; arrested oocytes are in panel B. All scale bars are 10 µm. See methods and text for description of thresholds used to categorize phase transitions. Statistical significance was determined using the Kruskal-Wallis test. **** indicates p<0.0001, *** indicates p<0.001, ** indicates p<0.01, * indicates p<0.05, ns indicates not significant. n=9-15.

We next asked if RNA binding protein phase transitions are induced as quickly as DAF-16 nuclear translocation. We first examined PGL-1, and found that in 30% of 11-20 min. worms, PGL-1 was at least partially decondensed out of granules, compared to the 0-10 min. worms which had bright PGL-1 granules as expected (Fig. 5B). We categorized PGL-1 as dispersed when the PGL-1 intensity in granules was below the lowest intensity observed in the 0-10 min. worms (see methods). The decondensation of PGL-1 was variable at all time points, and not statistically significant until the 51-60-min. worms; however, it was clear that at least modest decondensation occurs in a subset of worms within the same rapid time frame as DAF-16 nuclear translocation. We next assayed PGL-1 in arrested oocytes after 0-10 min. of imaging and observed a mix of small and large PGL-1 granules at the oocyte cortex as expected (Fig. 5C). PGL-1 appeared dispersed in 17% of worms at 11-20 min., and within 21-30 min. of imaging, we observed a significant increase in the percentage of worms with dispersed PGL-1 (Fig. 5C). At later time points, we continued to observe dispersal of PGL-1; however, the penetrance of the response was variable (Fig. 5C). These data suggest PGL-1 is not more stable when condensed into large granules in arrested oocytes; in fact, it appears more sensitive to imaging stress than when in small granules of diakinesis oocytes (Fig. 5B). Lastly, we examined the response of CAR-1 in diakinesis oocytes. In the 0-10 min. worms CAR-1 was mostly diffuse throughout the cytoplasm as expected (Fig. 5D). After 11-20 min., CAR-1 condensation was observed in 40% of worms; condensation was defined as worms having >3x as many CAR-1 granules at the cortex as the largest number of granules in 0-10 min. worms (Fig. 5D). At the three time points beyond 30 min., CAR-1 was condensed in a significant percent of worms (60-100%; Fig. 5D).

Based on these observations, we conclude that while imaging stress induces changes to both DAF-16 translocation and RNA binding protein condensation, the effects on RNA binding proteins are more variable. Nonetheless, inadvertent phase transitions of decondensation or condensation can occur within 11-20 min. of imaging which is fairly rapid, especially if researchers routinely prepare their worms at a location distant from their microscopy facility.

## Conclusions

The number of researchers studying *in vivo* phase-separated RNA binding proteins has grown tremendously over the past decade and resulted in a paradigm shift in our understanding of cellular organization. Here, we describe a potential pitfall of live-cell imaging using fluorescent reporter strains to study phase transitions. Our results demonstrate that liquid-like proteins, such as PGL-1, are extremely sensitive to decondensation during imaging, in as few as 11-20 minutes. Moreover, other RNA binding proteins are sensitive to ectopic condensation during imaging. Our results showing inadvertent phase transitions of germ line RNA binding proteins in the *C. elegans* model system seem likely to have broad applicability to live-cell imaging in all *in vivo* systems. Careful controls of imaging conditions are clearly critical to avoid inadvertent phase transitions. Furthermore, because the disruption of phase transitions can alter the function of RNA binding proteins, our results are pertinent not only when studying phase transitions, but for all physiological assays.

## ACKNOWLEDGEMENTS

We appreciate strains provided by Dr. Pam Padilla, Dr. Dustin Updike, Dr. Geraldine Seydoux, and the *Caenorhabditis* Genetics Center, which is funded by NIH Office of Research Infrastructure Programs (P40 OD010440). We thank Wormbase. C.M. was supported by an Undergraduate Summer Scholars grant from the Office of Research and Graduate Studies at Central Michigan University. Funding for this work came from a CMU Faculty Research and Creative Endeavors grant to J.S. and NIH grant 2R15GM109337-02A1 to J.S.

## Notes

### Competing Interest Statement

The authors have declared no competing interest.

### Summary of Updates

The revision includes the first four figures of the original, and the supplemental figure is now Figure 4. Prior figures 5 and 6 will be included in a separate manuscript.

